# Microbiota composition and evenness predict survival rate of oysters confronted to Pacific Oyster Mortality Syndrome

**DOI:** 10.1101/378125

**Authors:** Camille Clerissi, Julien de Lorgeril, Bruno Petton, Aude Lucasson, Jean-Michel Escoubas, Yannick Gueguen, Lionel Dégremont, Guillaume Mitta, Eve Toulza

**Author notes:** Corresponding author: Eve Toulza. Mailing address: IHPE UMR 5244, Université de Perpignan via Domitia, 52 Avenue Paul Alduy, F-66860 Perpignan Cedex, France., Camille Clerissi. Mailing address: CRIOBE USR 3278, Université de Perpignan via Domitia, 52 Avenue Paul Alduy, F-66860 Perpignan Cedex, France.

## Abstract

Pacific Oyster Mortality Syndrome (POMS) affects *Crassostrea gigas* oysters worldwide and caused important economic losses. Disease dynamics was recently deciphered and revealed a multiple and progressive infection caused by the *Ostreid herpesvirus* OsHV-1 µVar, triggering an immunosuppression followed by microbiota destabilization and bacteraemia by opportunistic bacterial pathogens. However, it remains unknown if microbiota might participate to oyster protection to POMS, and if microbiota characteristics might be predictive of oyster mortalities. To tackle this issue, we transferred full-sib progenies of resistant and susceptible oyster families from hatchery to the field during a period in favour of POMS. After five days of transplantation, oysters from each family were either sampled for individual microbiota analyses using 16S rRNA gene-metabarcoding or transferred into facilities to record their survival using controlled condition. As expected, all oysters from susceptible families died, and all oysters from the resistant family survived. Quantification of OsHV-1 and bacteria showed that five days of transplantation was long enough to contaminate oysters by POMS, but not for entering the pathogenesis process. Thus, it was possible to compare microbiota characteristics between resistant and susceptible oyster families at the early steps of infection. Strikingly, we found that microbiota evenness and abundances of Cyanobacteria (Subsection III, family I), Mycoplasmataceae, Rhodobacteraceae, and Rhodospirillaceae were significantly different between resistant and susceptible oyster families. We concluded that these microbiota characteristics might predict oyster mortalities.

## Introduction

The farmed oyster *Crassostrea gigas* is heavily affected by the Pacific Oyster Mortality Syndrome (POMS) targeting juveniles (Barbosa Solomieu et al., 2015; Pernet et al., 2016). This disease is multifactorial and depends on water temperature (Petton et al., 2015), development stage (Azéma et al., 2017), and oyster diet (Pernet et al., 2019). It is also polymicrobial due to the combined development of viral and bacterial infections (de Lorgeril et al., 2018). Different susceptibility levels were previously associated with oyster physiological status, genetic backgrounds (Dégremont et al., 2005; Samain et al., 2007; Wendling et al., 2017), in association with microbiota dysbiosis (Lokmer and Wegner, 2015).

Recently, holistic molecular approaches revealed the mechanism of POMS (de Lorgeril et al., 2018; Rubio et al., 2019). These studies showed that an infection by the Ostreid herpesvirus (OsHV-1 µVar) is the critical step in the infectious process leading to an immune-compromised state by altering hemocyte physiology. This first process is followed by a microbiota destabilization which “opens the door” to bacterial pathogens (*e.g.*, vibrios) that target hemocytes to induce their lysis. The infectious process is completed with subsequent bacteraemia, which is the ultimate step inducing oyster death.

So far, it is still unknown whether oyster microbial associates might influence disease development, and if microbiota characteristics might predict oyster mortalities. However, microbiota can play a role of physical barriers against pathogens. For example, it was suggested that part of the resident hemolymph bacteria may contribute to oyster protection by producing antimicrobial peptides (Desriac et al., 2014). Other studies highlighted protective effects of (i) secondary endosymbiont of aphids against parasitoid wasps (Oliver et al., 2003), (ii) toxic alkaloids produced by endophytic fungi of grasses against herbivores (Clay, 1990), or (iii) symbiotic bacteria of frogs against pathogenic fungi (Woodhams et al., 2007). Furthermore, microbial associates might also stimulate immunity of their hosts (Wang et al., 2019), and thus indirectly limit pathogen development.

To tackle this issue, we used one resistant oyster family (R_F21_) and two susceptible families (S_F15_ and S_F32_) used in a previous study (de Lorgeril et al., 2018). Pathogen-free oysters (reproduced and grown in bio-secured conditions) from these three full-sib families were placed for five days in the field during an infectious period (Figure 1). According to previous observations, 16°C was a relevant threshold to define the infectious period, as high mortality rates were observed above this temperature (Pernet et al., 2012; Petton et al., 2013; Dégremont et al., 2015). Moreover, five days were considered sufficient for oyster contamination by the causal agents of the disease (Petton et al., 2015), and allow microbiota analyses before disease development and animal death. In both the hatchery (control) and after five days of transplantation in the field, oysters were sampled, and we analyzed oyster-associated bacterial communities using 16S rRNA gene-metabarcoding.

**Figure 1.**
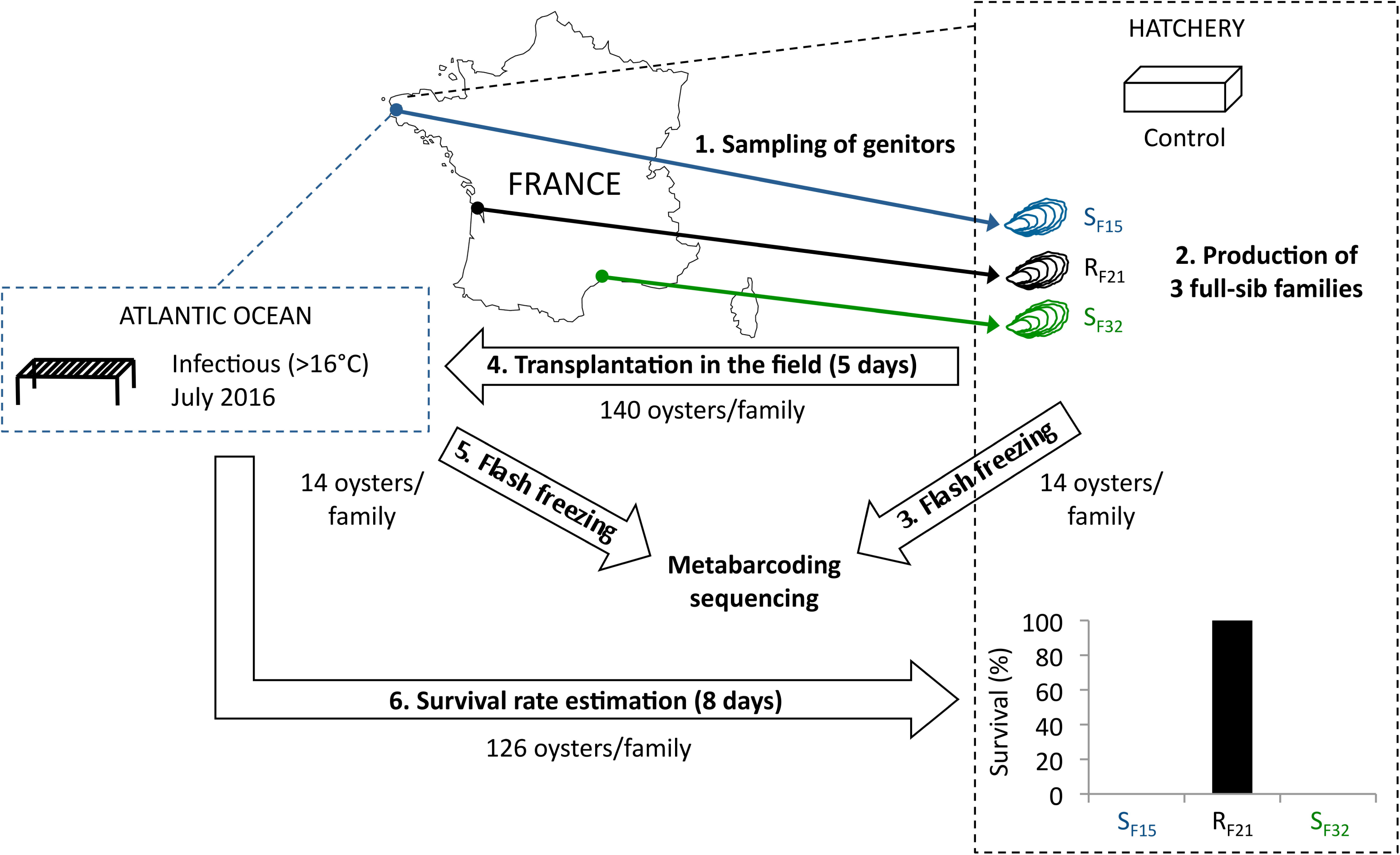
Experimental design. Three oyster full-sib families were produced in controlled condition (hatchery), and placed for five days in the environment (Atlantic Ocean, latitude: 48.335263; longitude: -4.317922) during an infectious period (July 2016). Then, oysters were flash frozen, DNA were extracted, and microbiota were sequenced using 16S rRNA gene-metabarcoding. Oysters deployed in the field during five days were followed during 8 days in the hatchery. Survival rates were recorded at endpoint.

This study aimed at comparing bacteria content of resistant and susceptible families in order to possibly identify microbiota features associated with oyster mortality and/or resistance. According to a previous holistic study (de Lorgeril et al., 2018), we expected that susceptible families had (i) early microbiota destabilization (increase of species richness, decrease of evenness, increase of microbiota dispersions), and (ii) possibly contained opportunistic and/or pathogenic bacteria (*Vibrio* sp., *Arcobacter* sp., etc.).

## Materials and Methods

### Full-sib *C. gigas* oyster families

In 2015, full-sib *C. gigas* oyster families were produced using a methodology that allowed the production of pathogen-free juveniles (de Lorgeril et al., 2018). Two oyster families (S_F15_ and S_F32_) were produced using one female and one male sampled from wild populations in the Atlantic Ocean and the Mediterranean Sea, respectively (Figure 1). In addition, family R_F21_ was produced using genitors from a mass selective breeding program aiming to increase the resistance of *C. gigas* oysters against OsHV-1. It was performed by breeding disease survivors throughout four generations of selection (Dégremont et al., 2015). Thus, each family (cohort) corresponded to the offspring of a biparental reproduction (full-sib progenies). Conditionning, reproduction and larval breeding were performed as described previously (de Lorgeril et al., 2018). At the larval and post-larval stages, oysters were fed with the same diet as the genitors at a concentration between 1500-2000 µm^3^ µl^−1^ (Rico-Villa et al., 2009). Before experiments, all oyster families were maintained in controlled condition at the laboratory (Argenton, France) using seawater treated with UV, filtered through 1 µm mesh, and enriched with a bi-specific phytoplankton diet made of *Tisochrysis lutea* (CCAP 927/14) and *Chaetoceros muelleri* (CCAP 1010/3) (in equal biomass proportion) at a ratio equivalent to 6% of the oyster dry mass (Rico-Villa et al., 2009). Finally, all oysters remained free of any abnormal mortality.

## Experimental design

About 140 juveniles per oyster family were either kept in the controlled condition or placed in the field for five days. The field site was located within a oyster farm in the Atlantic Ocean during an infectious period (Figure 1). This infectious period was selected according to seawater temperatures (above 16°C), and was confirmed by the observed mortality rates. After five days of transplantation (July 2016), no mortality occurred and 14 individuals per family were flash frozen in liquid nitrogen and stored at -80°C. The remaining oysters were then transferred into the hatchery under controlled conditions to monitor the survival rates of the three families. The number of dead oysters was recorded at day 13 (*i.e.*, eight days after the end of transplantation). Similarly, 14 individuals per family (except 13 for S_F32_) kept in the controlled condition were flash frozen in liquid nitrogen and stored at -80°C.

### DNA extraction, PCR and sequencing

Frozen oysters were ground in liquid nitrogen in 50 ml stainless steel bowls using 20 mm diameter grinding balls (Retsch MM400 mill). The powders were stored at -80°C, and were then used for DNA extractions using the DNA from tissue Macherey-Nagel kit (reference 740952.250) according to the manufacturer’s protocol. In order to improve DNA extractions, we added a crushing step, which consisted in an additional 12 minutes mechanical lysis using zirconium beads before the 90 min enzymatic lysis in the presence of proteinase K. DNA concentration and quality were checked with Epoch microplate spectrophotometer (BioTek Instruments, Inc.).

Then, the 16S rRNA gene of bacterial communities was amplified and sequenced using the variable V3V4 loops (341F: 5’-CCTACGGGNGGCWGCAG-3’; 805R: 5’-GACTACHVGGGTATCTAATCC-3’) (Klindworth et al., 2013). Paired-end sequencing (250 bp read length) was performed at the McGill University (Génome Québec Innovation Centre, Montréal, Canada) on the MiSeq system (Illumina) using the v2 chemistry according to the manufacturer’s protocol. Raw sequence data are available in the SRA database (BioProject ID PRJNA419907).

### Quantification of bacteria and OsHV-1

Quantification of OsHV-1, total bacteria 16S rDNA, and total *Vibrio* 16S rDNA was performed using quantitative PCR (qPCR). All amplification reactions were analysed using a Roche LightCycler 480 Real-Time thermocycler (qPHD-Montpellier GenomiX platform, Montpellier University, France). A Labcyte Acoustic Automated Liquid Handling Platform (ECHO) was used for pipetting into the 384-well plate (Roche). The total qPCR reaction volume was 1.5 µl and consisted of 0.5 µl DNA (40 ng µl-1) and 1 µl LightCycler 480 SYBR Green I Master mix (Roche) containing 0.5 µM PCR primer (Eurogenetec SA). Virus-specific primer pairs targeted a DNA polymerase catalytic subunit (DP, ORF100, AY509253): Fw-ATTGATGATGTGGATAATCTGTG and Rev-GGTAAATACCATTGGTCTTGTTCC (Davison et al., 2005). Total bacteria specific primer pairs were the 341F-CCTACGGGNGGCWGCAG and 805R-GACTACHVGGGTATCTAATCC primers targeting the variable V3V4 loops for bacterial communities (Klindworth et al., 2013). Total *Vibrio* specific primer pairs were Fw-GGCGTAAAGCGCATGCAGGT and Rev-GAAATTCTACCCCCCTCTACAG (Mansergh and Zehr, 2014). qPCR reactions were performed with the following program: 95 °C for 10 min, followed by 40 cycles of denaturation (95 °C, 10 s), hybridization (60 °C, 20 s) and elongation (72 °C, 25 s). After these PCR cycles a melting temperature curve of the amplicon was generated to verify the specificity of the amplification. Absolute quantification of OsHV-1 copies was calculated by comparing the observed Cq values to a standard curve generated from the DNA polymerase catalytic subunit amplification product cloned into the pCR4-TOPO vector. For total bacteria and total *Vibrio* 16S rDNA, we used the relative quantification calculated by the 2^−ΔΔCq^ method (Pfaffl, 2001) with the mean of the measured threshold cycle values of two reference genes (*Cg-BPI*, GenBank: AY165040, *Cg-BPI* F-ACGGTACAGAACGGATCTACG, *Cg-BPI* R-AATCGTGGCTGACATCGTAGC and *Cg-actin*, GenBank: AF026063, *Cg-actin* F-TCATTGCTCCACCTGAGAGG, *Cg-actin* R-AGCATTTCCTGTGGACAATGG).

### Sequence analyses

The FROGS pipeline (Find Rapidly OTU with Galaxy Solution) implemented into a galaxy instance (http://sigenae-workbench.toulouse.inra.fr/galaxy/) was used to define Operational Taxonomic Units (OTU), and computed taxonomic affiliations (Escudié et al., 2017). Briefly, paired reads were merged using FLASH (Magoc and Salzberg, 2011). After denoising and primer/adapters removal with cutadapt (Martin, 2011), *de novo* clustering was performed using SWARM that uses a local clustering threshold, with aggregation distance d=3 (Mahé et al., 2015). Chimera were removed using VSEARCH (*de novo* chimera detection) (Rognes et al., 2016). Particularly, this method divided each sequence into four fragments, and then looked for similarity with putative parents in the whole set of OTUs. We filtered the dataset for singletons and we annotated OTUs using Blast+ against the Silva database (release 123, September 2015) to produce an OTU and affiliation table in standard BIOM format. Rarefaction curves of species richness were produced using the {phyloseq} R package, and the ggrare function (McMurdie and Holmes, 2013). In order to compare samples for alpha and beta diversity, we used the rarefy_even_depth function to subsample dataset to 5148 reads per sample excluding chloroplasts. The alpha diversity metrics (Chao1 and Shannon) were estimated at the OTU level with the estimate_richness function. Moreover, Pielou’s measure of species evenness was computed using the diversity function in {vegan}. We also used phyloseq to obtain abundances at differents taxonomic ranks (for genus and family) (tax_glom function). Because agglomerate of multi-affiliation and unknown taxa does not make sense at higher taxonomic ranks, we only kept taxa having a true annotation for each corresponding taxonomic rank. We computed Bray-Curtis dissimilarities to study beta diversity, *i.e.*, distances between samples for OTU compositions (vegdist function, {vegan}).

### Statistical and multivariate analyses

All statistical analyses were done using R v3.3.1 (R: a language and environment for statistical computing, 2008; R Development Core Team, R Foundation for Statistical Computing, Vienna, Austria [http://www.R-project.org]).

Principal coordinate analyses (pcoa, {vegan}) were computed to describe compositions of microbial communities between samples using Bray-Curtis dissimilarities (vegdist, {vegan}). Multivariate homogeneity of group dispersions was tested between microbial assemblages of resistant and susceptible oyster families using 999 permutations (permutest, {vegan}). We used DESeq2 (Love et al., 2014) (DESeq {DESeq2}) to identify OTUs having differential abundances between resistant and susceptible oyster families. Heatmaps of significant bacterial genera and families were computed using relative abundances and the heatmap.2 function ({gplots}).

We performed one-way ANOVA or non-parametric Kruskal-Wallis tests (when normality of residuals was rejected (Shapiro test)) to compare alpha diversity metrics of microbiota. When ANOVA or Kruskal–Wallis tests were significant, we then computed pairwise comparisons between group levels (post-hoc analyses) using pairwise-t-tests or Dunn tests, respectively.

For all analyses, the threshold significance level was set at 0.05. P-values were corrected for multiple comparisons using Benjamini and Hochberg’s method (Benjamini and Hochberg, 1995) (p.adjust, {stats}).

## Results

### Survival rates and disease development

Full-sib progenies from one resistant (R_F21_) and two susceptible (S_F15_ and S_F32_) oyster families were produced and reared in bio-secured conditions (Argenton hatchery, France). About 140 pathogen-free oysters per family were transferred to the infectious environment for five days (Brest, Atlantic Ocean, temperature above 16°C) (Figure 1). Fourteen oysters per family were then flash frozen for individual bacterial microbiota analyses. The other 126 oysters per family were placed back in hatchery to monitor survival rates. As expected, all oysters from the two susceptible families died, whereas all oysters from the resistant family survived (Figure 1). This observation showed that oysters were contaminated during transplantation, and then developed the disease. Quantification of OsHV-1 and bacteria at day 5 post-transplantation showed that only three oysters from susceptible families displayed moderate to important viral infection, and that only one individual displayed both high viral infection and bacteraemia (Figure 2). These results suggested that five days of transplantation in the field during an infectious period were long enough for oyster contamination, but not for entering the pathogenesis process (except for three oysters).

**Figure 2.**
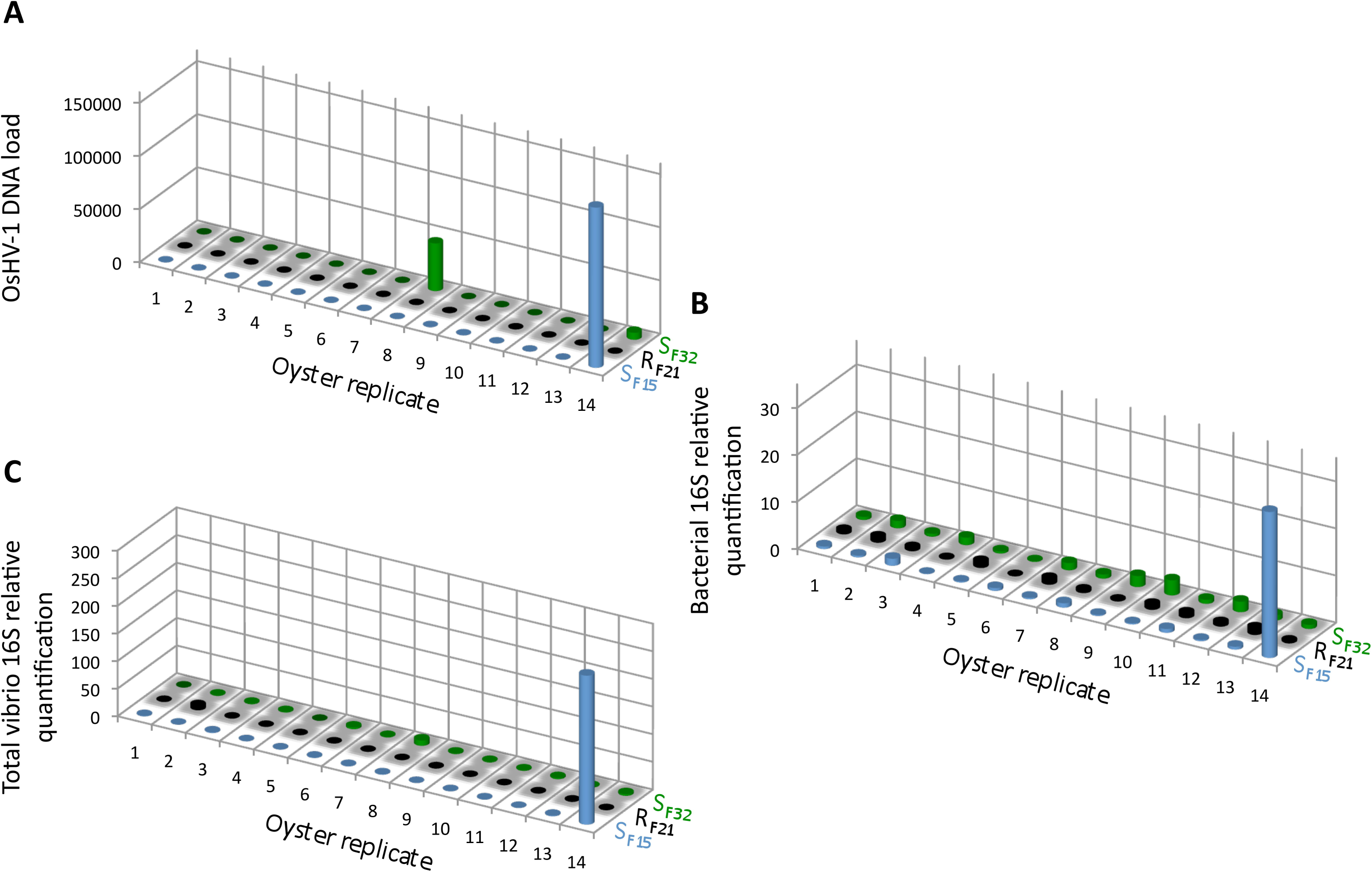
OsHV-1 and bacterial colonization in oysters at day 5 post-transplantation in field condition during an infectious period. **A.** The OsHV-1 load was quantified by qPCR and expressed as viral genomic units per ng of oyster DNA. Relative quantification of total bacteria (**B**), and total vibrio (**C**) abundance were measured by qPCR.

The whole bacterial communities were sequenced using the 16S rRNA gene from 14 oysters per family and per condition (except 13 oysters for S_F32_ in hatchery) (*i.e.*, 83 oyster-associated microbiota for control and infectious conditions). In average, each sample contained 21423 sequences representing 1075 OTUs (Supplementary Figure 1, Supplementary Tables 1 and 2). First, we searched for known opportunistic genera within microbiota of diseased oysters (de Lorgeril et al., 2018). Nine out of ten previously identified genera were present in our dataset (Figure 3). Most were abundant for the oyster highly infected by OsHV-1 and displaying bacteraemia (SF15.S.R14) (Figures 2 and 3A). Both *Vibrio* and *Psychromonas* genera were found within the three diseased oysters (Figure 3A). However, *Vibrio* occurred in the majority of healthy individuals as well (Figure 3B).

**Figure 3.**
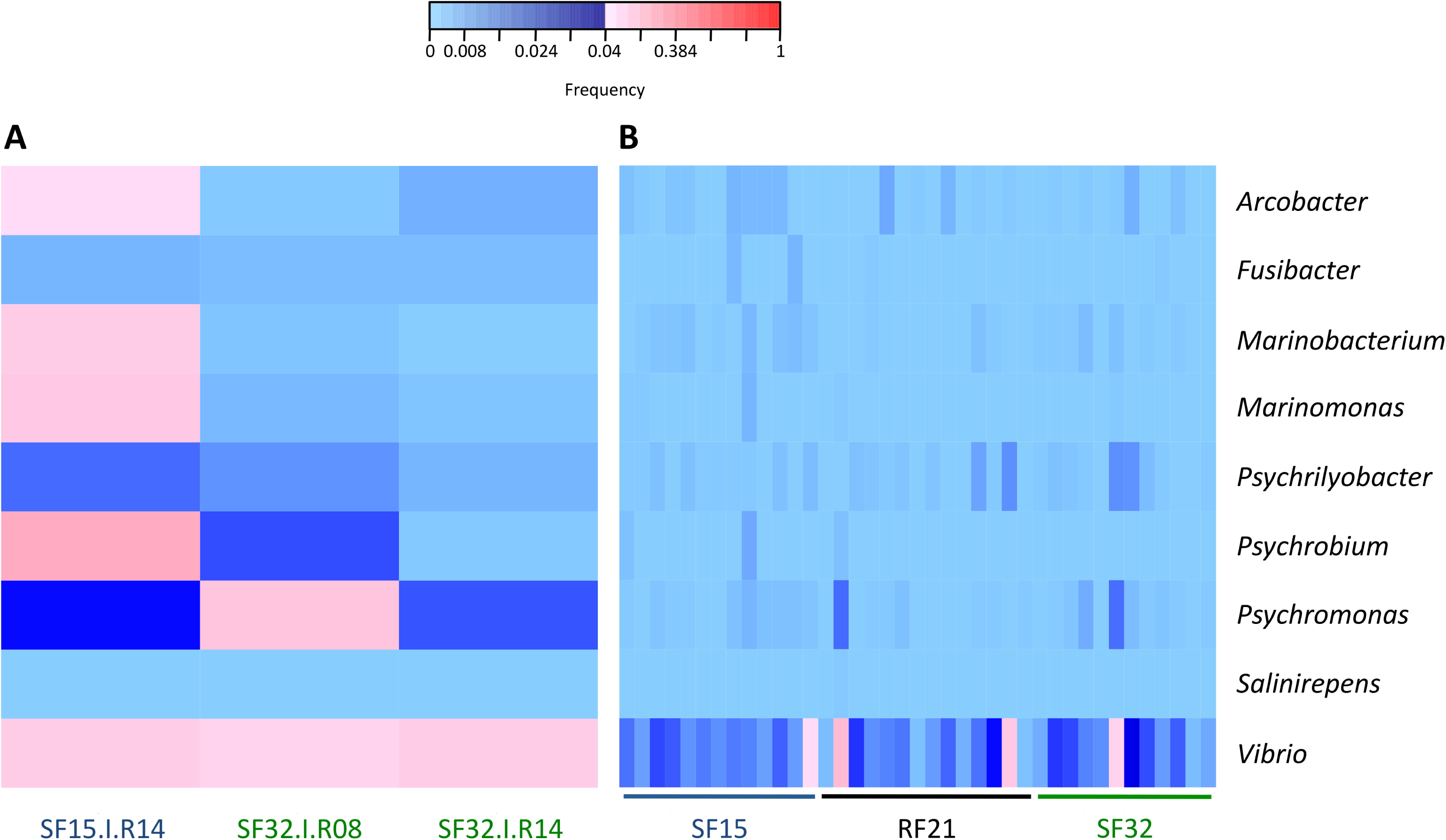
Frequencies of known opportunistic bacterial genera associated with POMS. **A.** Positive oysters for OsHV-1 detection (SF15.I.R14, SF32.I.R08, and SF32.I.R14). **B.** Negative oysters for OsHV-1 detection for each family. These opportunistic genera were identified in a previous experimental infection (de Lorgeril et al., 2018). Frequencies above and below 4% are displayed in red and blue, respectively.

### Microbiota assemblages before diseased development

Although five days of transplantation were long enough for oyster contamination, only three individuals entered the pathogenesis process as indicated by virus load. Thus, after discarding these three individuals, it was possible to compare microbiota characteristics of the other oysters between resistant and susceptible families before the onset of disease development.

First, we compared alpha diversity indices (Chao1, evenness and Shannon) between oyster families (Figure 4 and Supplementary Table 3). This analysis showed that Chao1 increased for all families between hatchery and the infectious environment. Moreover, evenness and Shannon indices increased only for the resistant family R_F21_. Notably, R_F21_ had higher evenness than S_F15_ and S_F32_ during the infectious condition. Secondly, we computed a principal coordinate analysis (PCoA) based on Bray–Curtis dissimilarities to describe microbiota compositions (Figure 5A). This ordination highlighted that oyster microbiota changed between hatchery and the infectious environment for the three oyster families. Unexpectedly, microbiota dispersion of R_F21_ was not lower than the susceptible families, and even showed higher dispersions for the infectious condition than in hatchery (Figure 5B).

**Figure 4.**
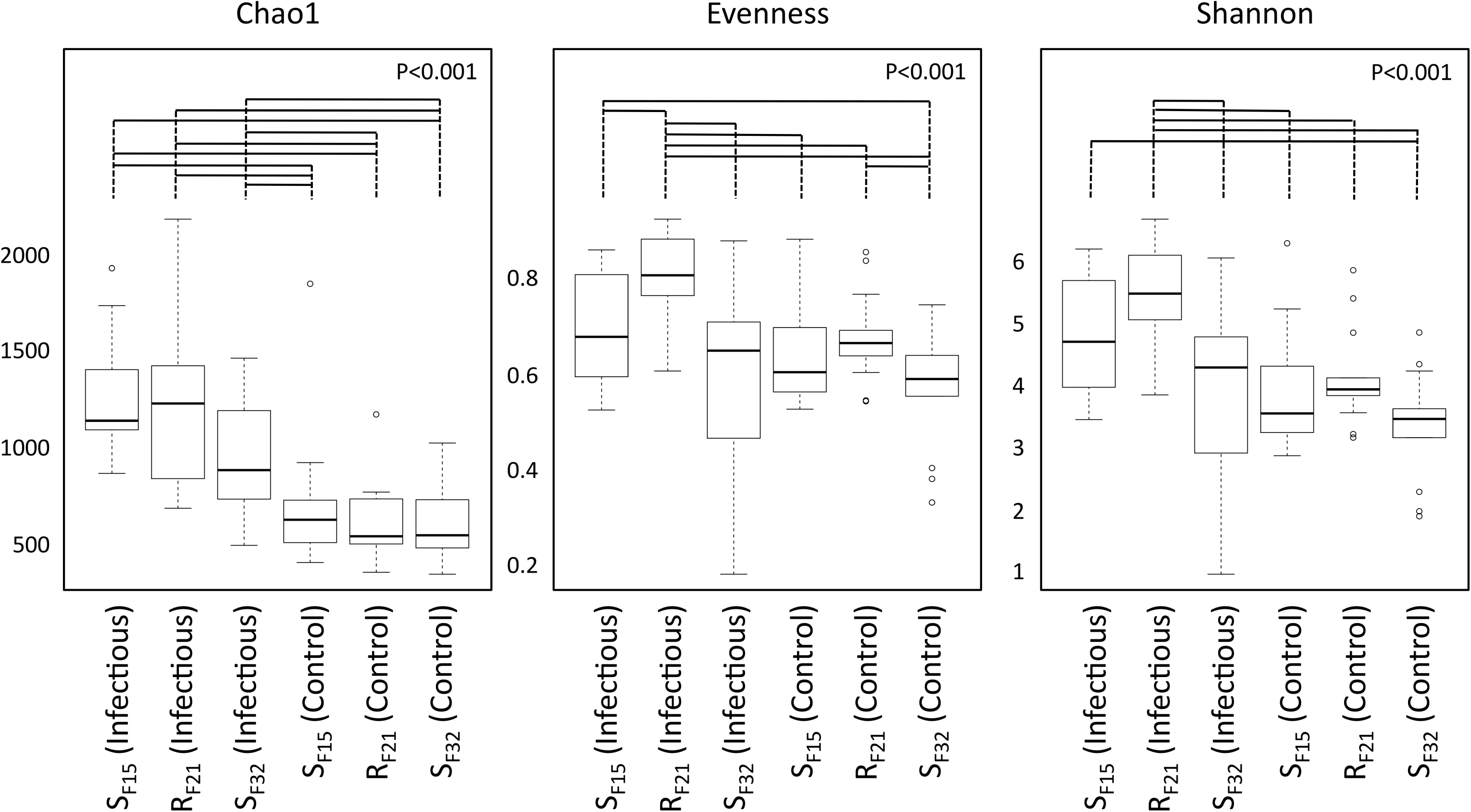
Comparison of alpha diversity indices between oyster families in controlled (hatchery) and infectious (field) conditions. P-values correspond to global tests. Horizontal lines above boxplots indicate significant pairwise comparisons between group levels (post-hoc analyses).

**Figure 5.**
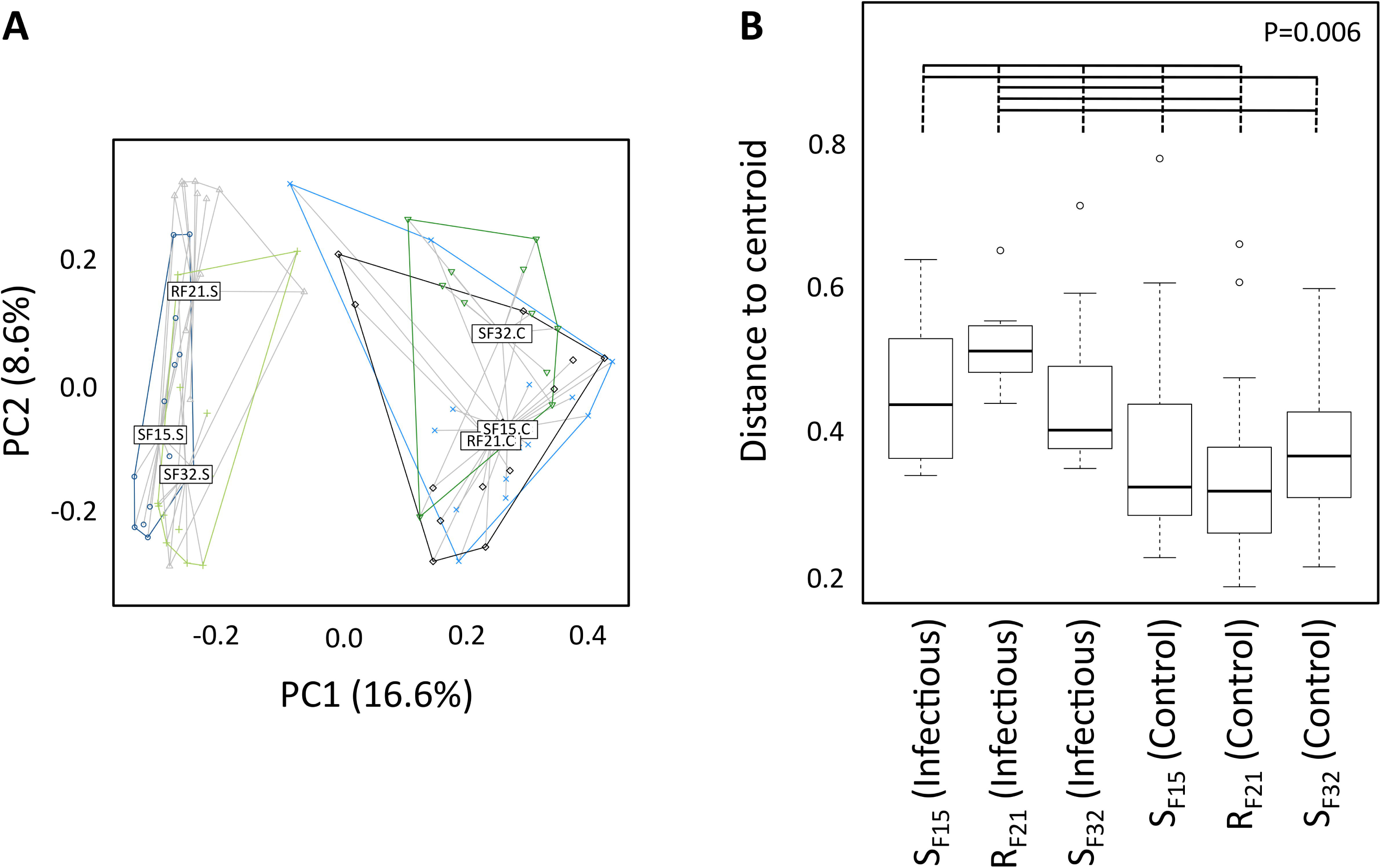
Microbiota compositions between oyster families in controlled (hatchery) and infectious (field) conditions. **A.** Principal coordinate analysis (Bray-Curtis dissimilarities between samples). **B.** Multivariate homogeneity of group dispersions. P-value corresponds to the global test. Horizontal lines above boxplots indicate significant pairwise comparisons between group levels (post-hoc analyses).

Altogether, these results suggested that microbiota assemblages highly changed between hatchery and the infectious environment for the three families. Although we did not expect to find higher microbiota dispersion for R_F21_, our analyses revealed that the resistant family had higher evenness than the susceptible families before the onset of disease development.

### Different bacteria between oyster families in hatchery and infectious environment

Because resistant and susceptible oyster families had different microbiota characteristics (evenness and dispersion) at the early steps of infection, we compared abundances of bacterial taxa within hatchery to identify putative pathogens, opportunists or mutualists that were consistently presents in microbiota before transplantation. Abundances were compared at two taxonomic ranks, bacterial genera (Supplementary Table 4) and families (Supplementary Table 5). Two genera (*Colwellia* and *Photobacterium*) (Supplementary Figure 2) and four families (Anaplasmataceae, Colwelliaceae, Mycoplasmataceae, and Vibrionaceae) (Figure 6) showed high abundances (>4% in at least one sample) and were significantly different between resistant and susceptible oyster families.

**Figure 6.**
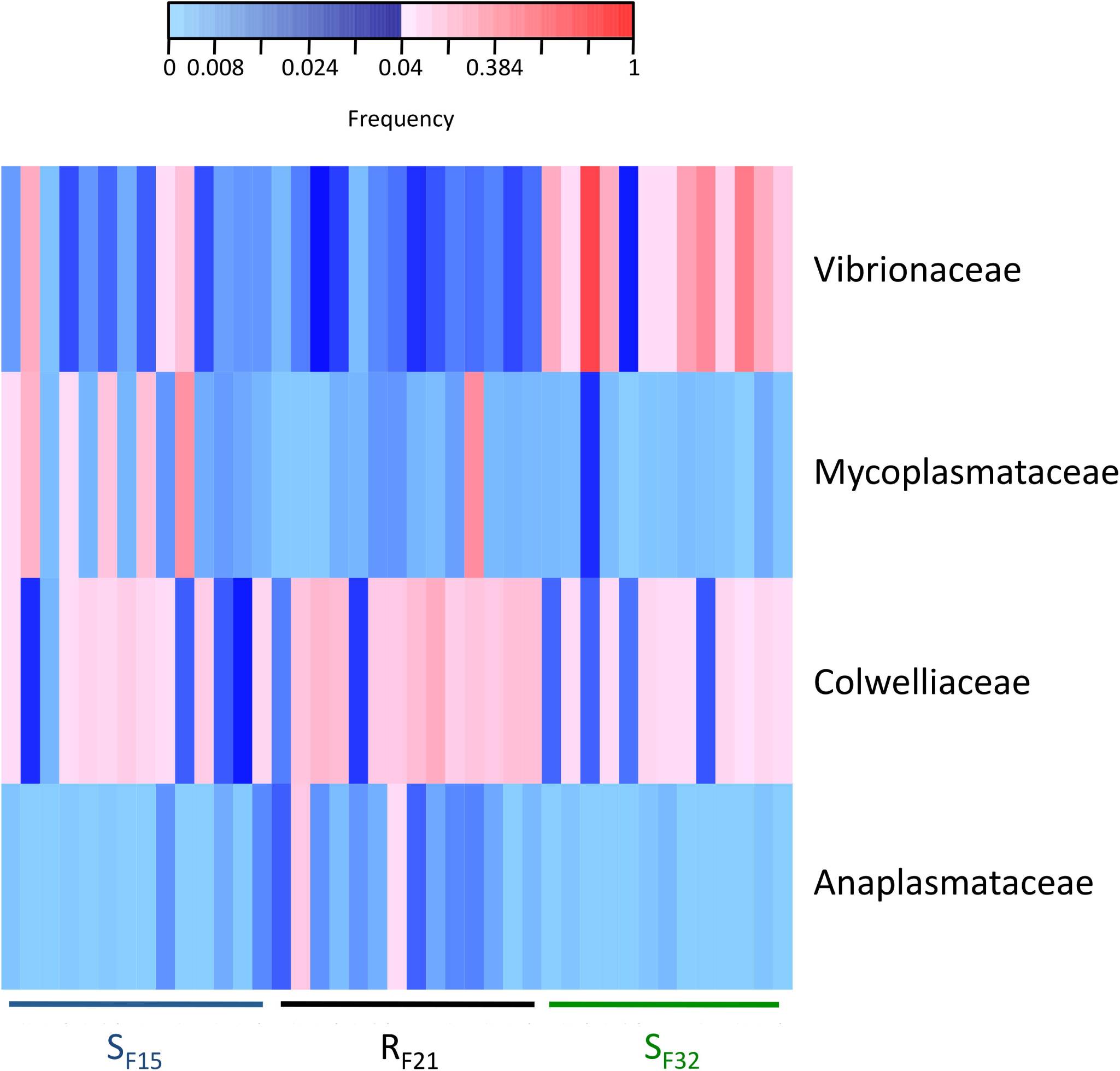
Heatmap of bacterial families that were significantly different between resistant and susceptible oyster families in the controlled condition (hatchery). Only bacterial families with a frequency above 4% in at least one sample are shown. Frequencies above and below 4% are displayed in red and blue, respectively.

We also compared bacterial taxa before disease development between resistant and susceptible oyster families that were placed in the infectious environment. Except for Mycoplasmataceae, significant bacterial taxa were not similar to hatchery-identified candidates (Figure 7 and Supplementary Figure 3). Four genera (*Phormidium, Pseudomonas, Thalassospira*, and *Umboniibacter*) (Supplementary Figure 3) and seven families (Cyanobacteria (Subsection I, family I), Cyanobacteria (Subsection III, family I), Mycoplasmataceae, Pseudomonadaceae, Rhodobacteraceae, Rhodospirillaceae, and Spirochaetaceae) (Figure 7) had high abundances (>4% in at least one sample) and were significantly different between resistant and susceptible oyster families.

**Figure 7.**
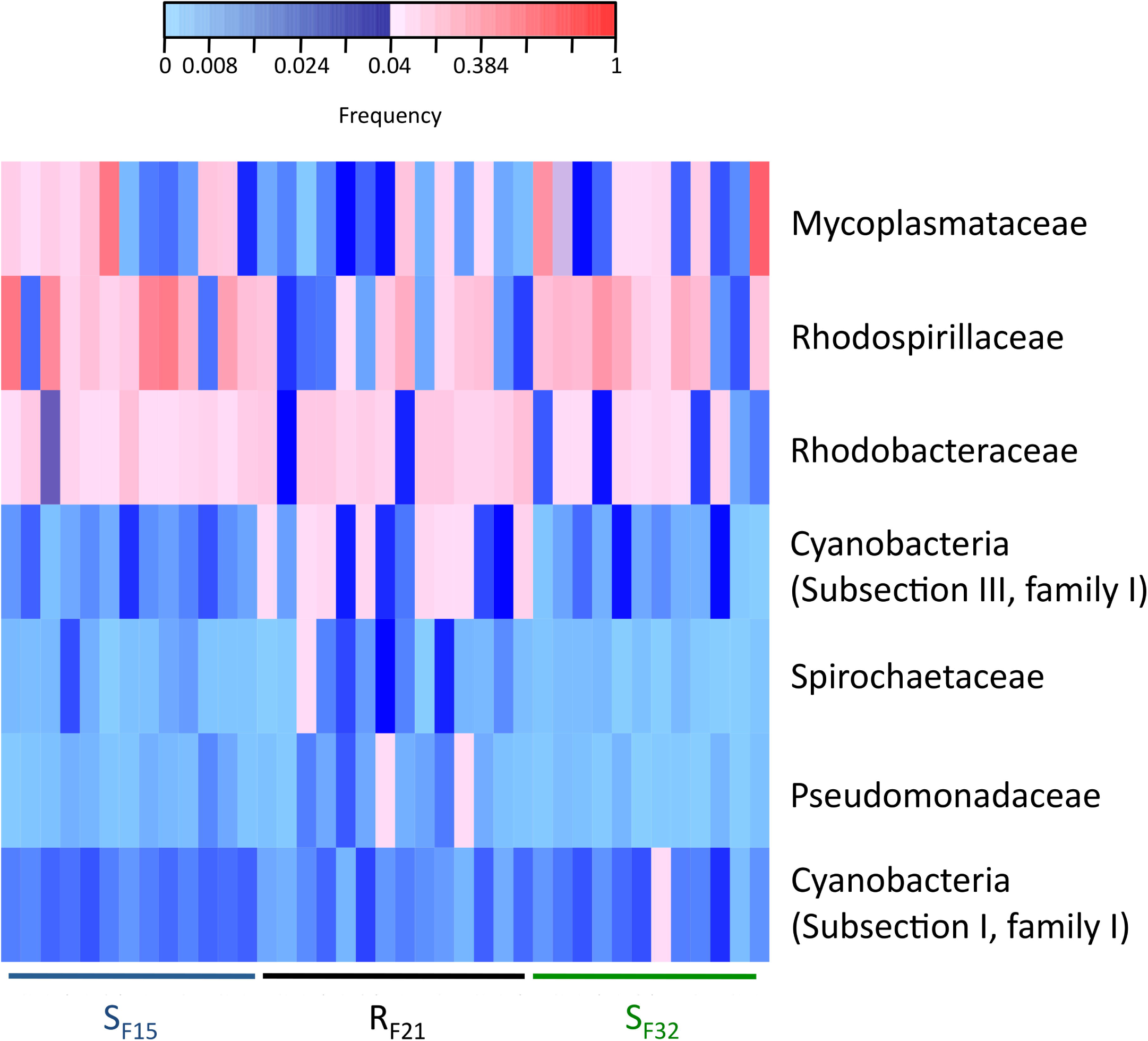
Heatmap of bacterial families that were significantly different between resistant and susceptible oyster families in the infectious condition (field). Only bacterial families with a frequency above 4% in at least one sample are shown. Frequencies above and below 4% are displayed in red and blue, respectively.

## Discussion

### Five days of transplantation allowed oyster contamination, but not disease development

Oysters were transplanted for five days in the field during an infectious period. According to previous observations (Dégremont, 2011; Petton et al., 2015), this time period was considered sufficient for oyster contamination. We did not observed dead oysters when they were sampled, but all individuals of susceptible families died eight days after they were placed back to laboratory tanks. This observation showed that oysters were contaminated during transplantation, and this exposure was sufficient for the further development of the complete pathogenesis process.

High abundances of OsHV-1 and total bacteria were previously observed for a susceptible family during disease development (de Lorgeril et al., 2018). In this study, most oysters (except three) did not exhibit high abundances of OsHV-1 or bacteria after five days of transplantation. As a consequence and after exclusion of these three oysters, we thus analysed oyster microbiota before the onset of disease development (*i.e.*, at the early steps of infection).

### Microbiota evenness was linked to oyster resistance during transplantation

According to a previous study (de Lorgeril et al., 2018), we expected to observe many modifications of microbiota for susceptible oyster families during transplantation, but not or few for the resistant one. In particular, we expected increase of alpha diversity indices (Chao1 and Shannon), and of microbiota dispersion for the susceptible families. Furthermore, we hypothesized that the resistant family might have stable microbiota, because this characteristic was linked to host homeostasis in many studies (Faith et al., 2013; Rungrassamee et al., 2016; Zaneveld et al., 2017).

Unexpectedly, we found that alpha diversity indices and dispersion of microbiota highly changed for the three oyster families (susceptible as well as resistant) when they were transferred from hatchery to the infectious environment. Microbiota evenness was the only index that discriminated resistant and susceptible families. It highlighted that OTU abundances within resistant oysters were more equally distributed than within susceptible oysters. This index may possibly have important effects on oyster homeostasis, because it was already found to be positively linked to ecosystem productivity (Wilsey and Potvin, 2000; Zhang et al., 2012), functional stability (Balvanera et al., 2005; Wittebolle et al., 2009), and invasion resistance (Wilsey and Polley, 2002; De Roy et al., 2013).

### Putative opportunistic and/or pathogenic bacteria of susceptible oysters

To date, most *C. gigas* diseases affecting juveniles in Europe were associated with *herpesvirus* (OsHV-1 µVar) and *Vibrio* strains (Labreuche et al., 2006; Garcia et al., 2011; Travers et al., 2014; Batista et al., 2015; Petton et al., 2015). Furthermore, other studies highlighted the role of *Roseovarius crassostreae* (Boettcher, 2005; Maloy et al., 2007), *Tenacibaculum soleae* (Burioli et al., 2017), and possibly Chlamydiales (Renault and Cochennec, 1995) and *Arcobacter* sp. (Lokmer and Wegner, 2015; Burioli et al., 2017). Other potential opportunistic bacteria were recently identified (de Lorgeril et al., 2018), such as *Cryomorphaceae, Fusibacter, Marinobacterium, Marinomonas, Psychrilyobacter, Psychrobium, Psychromonas*, and *Salinirepens*.

In this study, most of these genera were identified within the oyster that displayed high abundances of OsHV-1 and bacteraemia. *Psychromonas* occurred within the three diseased oysters, and might possibly be involved early in disease development, even if its pathogenic role was not demonstrated so far. Furthermore, three bacterial families (Mycoplasmataceae, Rhodospirillaceae, and Vibrionaceae) were linked to susceptible oysters in hatchery and/or before disease development in the field. Among Vibrionaceae, *Photobacterium* genus was mostly associated with S_F32_ in hatchery, and as well with low disease resistance oysters in a previous study (King et al., 2019). Among Rhodospirillaceae, *Thalassospira* genus was already identified in *C. gigas* oysters (Fernandez-Piquer et al., 2012). Even though Rhodospirillaceae were found in diseased tissues of *Platygyra carnosus* corals (Ng et al., 2015), there is no evidence that they may act as pathogens for oysters until now. Lastly, Mycoplasmataceae family was abundant in *C. virginica* oysters from Hackberry Bay (King et al., 2012), and decreased in oysters during experimental infections (de Lorgeril et al., 2018). However, it was intriguing to find in this study that Mycoplasmataceae abundances were significantly lower for resistant than susceptible oyster families in both hatchery and field conditions, at the early steps of infection.

### Putative beneficial bacteria of resistant oysters

Some observations suggested that bacteria might protect oysters from pathogens. For example, *Aeromonas media* A199, *Phaeobacter gallaeciensis* and *Pseudoalteromonas* sp. D41 improved survival of *C. gigas* against *Vibrio* strains when they were tested as probiotics (Gibson et al., 1998; Kesarcodi-Watson et al., 2012). In particular, the *Pseudoalteromonas* genus is known to produce a wide variety of biologically active secondary metabolites (Kalinovskaya et al., 2004; Bowman, 2007). Finding beneficial bacteria could be interesting and pave the way for prophylactic measures in oyster farming.

Here we identified three bacterial families associated with resistant oysters: Colwelliaceae, Cyanobacteria (Subsection III, family I), and Rhodobacteraceae. Among Colwelliaceae, the *Colwellia* genus was already found within *C. gigas* microbiota (Madigan et al., 2014). In this study, this family and genus had high abundances for resistant oysters in hatchery, but they were not significantly different between resistant and susceptible oyster families during transplantation, suggesting a limited role in POMS. Moreover, although Rhodobacteraceae was associated to juvenile oyster disease (Boettcher, 2005), this bacterial family was commonly associated with oysters (Asmani et al., 2016; Takenaka et al., 2017; Arahal et al., 2018). Strikingly, Cyanobacteria (Subsection III, family I) had low abundances in all susceptible oyster microbiota before disease development, and were abundant in most resistant oysters. Although cyanobacteria had a negative effect on other marine invertebrates such as scleractinian corals (Meyer et al., 2016), they were already observed at high relative abundances within oyster microbiota (Chauhan et al., 2014). Particularly, they were abundant in the digestive gland, connective tissue, mantle, and gonad of oysters (Avila-Poveda et al., 2014). Because cyanobacteria persisted in oyster tissues without signs of alterations, a possible endosymbiotic relationship was even proposed (Avila-Poveda et al., 2014). Notably, cyanobacteria were also negatively correlated to culturable *Vibrio* abundance in free-living microbial communities (Turner et al., 2009). Hence, cyanobacteria might play a role of barrier against pathogens from the genus *Vibrio*. In particular, photosynthetic activity of cyanobacteria could inhibited pathogen growth through the production of reactive oxygen species, such as observed for *Plasmodium* infection in *Anopheles gambiae* (Cirimotich et al., 2011).

To conclude, we placed oysters from resistant and susceptible families in field condition for five days during a period in favour of POMS. The analyses of oyster microbiota at the early steps of infection revealed differences between resistant and susceptible oyster families. These differences suggested that both the structure (evenness) and the composition (putative opportunists, pathogens, mutualists) might predict oyster mortalities. Future studies should test the role of Cyanobacteria (Subsection III, family I), Mycoplasmataceae, Rhodobacteraceae, and Rhodospirillaceae within *C. gigas* microbiota. In particular, these studies should evaluate the possibility of using Cyanobacteria (Subsection III, family I) as probiotics for oyster farming.

## Declarations

### Ethics approval and consent to participate

Not applicable.

### Consent for publication

Not applicable.

### Availability of data and materials

The datasets generated during the current study are available in the Sequence Read Archive repository under BioProject ID PRJNA419907 (to be released upon publication).

### Competing interests

The authors declare that they have no competing interests.

### Funding

CC benefited of post-doctoral fellowships from CNRS and IFREMER. This work was supported by the French National Research Agency ANR, project ANR-14-CE19-0023 DECIPHER (coordinator G. Mitta) and the DHOF program of the UMR5244/IHPE (http://ihpe.univ-perp.fr/en/ihpe-transversal-holobiont/). This project has received funding from the European Union’s Horizon 2020 Research and innovation programme under grant agreement N° 678589 (VIVALDI project).

### Authors’ contributions

CC, JDL, JME, YG, LD, GM and ET were involved in the study concept and design. BP was involved in the generation and maintaining of all animals used in this study. JDL, BP, JME, YG, GM and ET were involved in the collection of samples. CC, JDL, AL and ET were involved in data acquisition and analysis. CC and ET drafted the manuscript and all authors contributed to critical revisions and approved the final manuscript.

## Acknowledgements

We thank the staff of the Ifremer stations of Argenton (LPI, PFOM) and Sète (LER), and the Comité Régional de Conchyliculture de Méditerranée (CRCM) for technical support in the collection of the oyster genitors and reproduction. We are grateful to IHPE members for stimulating discussions. We thank the genotoul bioinformatics platform Toulouse Midi-Pyrenees and Sigenae group for providing help and computing resources thanks to Galaxy instance http://sigenae-workbench.toulouse.inra.fr.

## References

Arahal, D. R., Lucena, T., Rodrigo-Torres, L., and Pujalte, M. J. (2018). *Ruegeria denitrificans* sp. nov., a marine bacterium in the family Rhodobacteraceae with the potential ability for cyanophycin synthesis. Int. J. Syst. Evol. Microbiol. 68, 2515–2522. doi:10.1099/ijsem.0.002867.

Asmani, K., Petton, B., Le Grand, J., Mounier, J., Robert, R., and Nicolas, J.-L. (2016). Establishment of microbiota in larval culture of Pacific oyster, *Crassostrea gigas*. Aquaculture 464, 434–444. doi:10.1016/j.aquaculture.2016.07.020.

Avila-Poveda, O. H., Torres-Ariño, A., Girón-Cruz, D. A., and Cuevas-Aguirre, A. (2014). Evidence for accumulation of *Synechococcus elongatus* (Cyanobacteria: Cyanophyceae) in the tissues of the oyster *Crassostrea gigas* (Mollusca: Bivalvia). Tissue Cell 46, 379–387. doi:10.1016/j.tice.2014.07.001.

Azéma, P., Lamy, J.-B., Boudry, P., Renault, T., Travers, M.-A., and Dégremont, L. (2017). Genetic parameters of resistance to *Vibrio aestuarianus*, and OsHV-1 infections in the Pacific oyster, *Crassostrea gigas*, at three different life stages. Genet. Sel. Evol. 49, 23. doi:10.1186/s12711-017-0297-2.

Balvanera, P., Kremen, C., and Martínez-Ramos, M. (2005). Applying community structure analysis to ecosystem function: examples from pollination and carbon storage. Ecol. Appl. 15, 360–375.

Barbosa Solomieu, V., Renault, T., and Travers, M.-A. (2015). Mass mortality in bivalves and the intricate case of the Pacific oyster, *Crassostrea gigas*. J. Invertebr. Pathol. 131, 2–10. doi:10.1016/j.jip.2015.07.011.

Batista, F. M., López-Sanmartín, M., Grade, A., Morgado, I., Valente, M., Navas, J. I., et al. (2015). Sequence variation in ostreid herpesvirus 1 microvar isolates detected in dying and asymptomatic *Crassostrea angulata* adults in the Iberian Peninsula: insights into viral origin and spread. Aquaculture 435, 43–51. doi:10.1016/j.aquaculture.2014.09.016.

Benjamini, Y., and Hochberg, Y. (1995). Controlling the false discovery rate: a practical and powerful approach to multiple testing. J. R. Stat. Soc. Ser. B, 289–300.

Boettcher, K. J. (2005). *Roseovarius crassostreae* sp. nov., a member of the *Roseobacter* clade and the apparent cause of juvenile oyster disease (JOD) in cultured Eastern oysters. Int. J. Syst. Evol. Microbiol. 55, 1531–1537. doi:10.1099/ijs.0.63620-0.

Bowman, J. P. (2007). Bioactive compound synthetic capacity and ecological significance of marine bacterial genus *Pseudoalteromonas*. Mar Drugs, 22.

Burioli, E. A. V., Varello, K., Trancart, S., Bozzetta, E., Gorla, A., Prearo, M., et al. (2017). First description of a mortality event in adult Pacific oysters in Italy associated with infection by a *Tenacibaculum soleae* strain. J. Fish Dis., 1–7 doi:10.1111/jfd.12698.

Chauhan, A., Wafula, D., Lewis, D. E., and Pathak, A. (2014). Metagenomic assessment of the Eastern oyster-associated microbiota. Genome Announc. 2, e01083-14-e01083-14. doi:10.1128/genomeA.01083-14.

Cirimotich, C. M., Dong, Y., Clayton, A. M., Sandiford, S. L., Souza-Neto, J. A., Mulenga, M., et al. (2011). Natural microbe-mediated refractoriness to plasmodium infection in *Anopheles gambiae*. Science 332, 855–858. doi:10.1126/science.1201618.

Clay, K. (1990). Fungal endophytes of grasses. Annu. Rev. Ecol. Syst. 21, 275–297.

Davison, A. J., Trus, B. L., Cheng, N., Steven, A. C., Watson, M. S., Cunningham, C., et al. (2005). A novel class of herpesvirus with bivalve hosts. J. Gen. Virol. 86, 41–53. doi:10.1099/vir.0.80382-0.

de Lorgeril, J., Lucasson, A., Petton, B., Toulza, E., Montagnani, C., Clerissi, C., et al. (2018). Immune-suppression by OsHV-1 viral infection causes fatal bacteraemia in Pacific oysters. Nat. Commun. 9, 4215. doi:10.1038/s41467-018-06659-3.

De Roy, K., Marzorati, M., Negroni, A., Thas, O., Balloi, A., Fava, F., et al. (2013). Environmental conditions and community evenness determine the outcome of biological invasion. Nat. Commun. 4, 1383. doi:10.1038/ncomms2392.

Dégremont, L. (2011). Evidence of herpesvirus (OsHV-1) resistance in juvenile *Crassostrea gigas* selected for high resistance to the summer mortality phenomenon. Aquaculture 317, 94–98. doi:10.1016/j.aquaculture.2011.04.029.

Dégremont, L., Bédier, E., Soletchnik, P., Ropert, M., Huvet, A., Moal, J., et al. (2005). Relative importance of family, site, and field placement timing on survival, growth, and yield of hatchery-produced Pacific oyster spat (*Crassostrea gigas*). Aquaculture 249, 213–229. doi:10.1016/j.aquaculture.2005.03.046.

Dégremont, L., Nourry, M., and Maurouard, E. (2015). Mass selection for survival and resistance to OsHV-1 infection in *Crassostrea gigas* spat in field conditions: response to selection after four generations. Aquaculture 446, 111–121. doi:10.1016/j.aquaculture.2015.04.029.

Desriac, F., Le Chevalier, P., Brillet, B., Leguerinel, I., Thuillier, B., Paillard, C., et al. (2014). Exploring the hologenome concept in marine bivalvia: haemolymph microbiota as a pertinent source of probiotics for aquaculture. FEMS Microbiol. Lett. 350, 107–116. doi:10.1111/1574-6968.12308.

Escudié, F., Auer, L., Bernard, M., Mariadassou, M., Cauquil, L., Vidal, K., et al. (2017). FROGS: Find, Rapidly, OTUs with Galaxy Solution. Bioinformatics doi:10.1093/bioinformatics/btx791. doi:10.1093/bioinformatics/btx791.

Faith, J. J., Guruge, J. L., Charbonneau, M., Subramanian, S., Seedorf, H., Goodman, A. L., et al. (2013). The long-term stability of the human gut microbiota. Science 341, 1237439–1237439. doi:10.1126/science.1237439.

Fernandez-Piquer, J., Bowman, J. P., Ross, T., and Tamplin, M. L. (2012). Molecular analysis of the bacterial communities in the live Pacific oyster (*Crassostrea gigas*) and the influence of postharvest temperature on its structure. J. Appl. Microbiol. 112, 1134–1143. doi:10.1111/j.1365-2672.2012.05287.x.

Garcia, C., Thébault, A., Dégremont, L., Arzul, I., Miossec, L., Robert, M., et al. (2011). Ostreid herpesvirus 1 detection and relationship with *Crassostrea gigas* spat mortality in France between 1998 and 2006. Vet. Res. 42, 73. doi:10.1186/1297-9716-42-73.

Gibson, L. F., Woodworth, J., and George, A. M. (1998). Probiotic activity of *Aeromonas media* on the Pacific oyster, *Crassostrea gigas*, when challenged with *Vibrio tubiashii*. Aquaculture 169, 111–120. doi:10.1016/S0044-8486(98)00369-X.

Kalinovskaya, N. I., Ivanova, E. P., Alexeeva, Y. V., Gorshkova, N. M., Kuznetsova, T. A., Dmitrenok, A. S., et al. (2004). Low-molecular-weight, biologically active compounds from marine *Pseudoalteromonas* species. Curr. Microbiol. 48, 441–446. doi:10.1007/s00284-003-4240-0.

Kesarcodi-Watson, A., Miner, P., Nicolas, J.-L., and Robert, R. (2012). Protective effect of four potential probiotics against pathogen-challenge of the larvae of three bivalves: Pacific oyster (*Crassostrea gigas*), flat oyster (*Ostrea edulis*) and scallop (*Pecten maximus*). Aquaculture 344–349, 29–34. doi:10.1016/j.aquaculture.2012.02.029.

King, G. M., Judd, C., Kuske, C. R., and Smith, C. (2012). Analysis of stomach and gut microbiomes of the Eastern oyster (*Crassostrea virginica*) from coastal Louisiana, USA. PLoS ONE 7, e51475. doi:10.1371/journal.pone.0051475.

King, W. L., Siboni, N., Williams, N. L. R., Kahlke, T., Nguyen, K. V., Jenkins, C., et al. (2019). Variability in the composition of Pacific oyster microbiomes across oyster families exhibiting different levels of susceptibility to OsHV-1 µvar disease. Front. Microbiol. 10, 473. doi:10.3389/fmicb.2019.00473.

Klindworth, A., Pruesse, E., Schweer, T., Peplies, J., Quast, C., Horn, M., et al. (2013). Evaluation of general 16S ribosomal RNA gene PCR primers for classical and next-generation sequencing-based diversity studies. Nucleic Acids Res. 41, e1. doi:10.1093/nar/gks808.

Labreuche, Y., Soudant, P., Goncalves, M., Lambert, C., and Nicolas, J. (2006). Effects of extracellular products from the pathogenic *Vibrio aestuarianus* strain 01/32 on lethality and cellular immune responses of the oyster *Crassostrea gigas*. Dev. Comp. Immunol. 30, 367–379. doi:10.1016/j.dci.2005.05.003.

Lokmer, A., and Wegner, K. M. (2015). Hemolymph microbiome of Pacific oysters in response to temperature, temperature stress and infection. ISME J. 9, 670–682.

Love, M. I., Huber, W., and Anders, S. (2014). Moderated estimation of fold change and dispersion for RNA-seq data with DESeq2. Genome Biol. 15, 550. doi:10.1186/s13059-014-0550-8.

Madigan, T. L., Bott, N. J., Torok, V. A., Percy, N. J., Carragher, J. F., de Barros Lopes, M. A., et al. (2014). A microbial spoilage profile of half shell Pacific oysters (*Crassostrea gigas*) and Sydney rock oysters (*Saccostrea glomerata*). Food Microbiol. 38, 219–227. doi:10.1016/j.fm.2013.09.005.

Magoc, T., and Salzberg, S. L. (2011). FLASH: fast length adjustment of short reads to improve genome assemblies. Bioinformatics 27, 2957–2963. doi:10.1093/bioinformatics/btr507.

Mahé, F., Rognes, T., Quince, C., de Vargas, C., and Dunthorn, M. (2015). Swarm v2: highly-scalable and high-resolution amplicon clustering. PeerJ 3, e1420. doi:10.7717/peerj.1420.

Maloy, A. P., Ford, S. E., Karney, R. C., and Boettcher, K. J. (2007). *Roseovarius crassostreae*, the etiological agent of Juvenile Oyster Disease (now to be known as Roseovarius Oyster Disease) in *Crassostrea virginica*. Aquaculture 269, 71–83. doi:10.1016/j.aquaculture.2007.04.008.

Mansergh, S., and Zehr, J. P. (2014). *Vibrio* diversity and dynamics in the Monterey Bay upwelling region. Front. Microbiol. 5, 48. doi:10.3389/fmicb.2014.00048.

Martin, M. (2011). Cutadapt removes adapter sequences from high-throughput sequencing reads. EMBnet J. 17, pp–10.

McMurdie, P. J., and Holmes, S. (2013). phyloseq: An R package for reproducible interactive analysis and graphics of microbiome census data. PLoS ONE 8, e61217. doi:10.1371/journal.pone.0061217.

Meyer, J. L., Gunasekera, S. P., Scott, R. M., Paul, V. J., and Teplitski, M. (2016). Microbiome shifts and the inhibition of quorum sensing by Black Band Disease cyanobacteria. ISME J. 10, 1204–1216.

Ng, J. C. Y., Chan, Y., Tun, H. M., Leung, F. C. C., Shin, P. K. S., and Chiu, J. M. Y. (2015). Pyrosequencing of the bacteria associated with *Platygyra carnosus* corals with skeletal growth anomalies reveals differences in bacterial community composition in apparently healthy and diseased tissues. Front. Microbiol. 6, 1142. doi:10.3389/fmicb.2015.01142.

Oliver, K. M., Russell, J. A., Moran, N. A., and Hunter, M. S. (2003). Facultative bacterial symbionts in aphids confer resistance to parasitic wasps. Proc. Natl. Acad. Sci. 100, 1803–1807.

Pernet, F., Barret, J., Le Gall, P., Corporeau, C., Dégremont, L., Lagarde, F., et al. (2012). Mass mortalities of Pacific oysters *Crassostrea gigas* reflect infectious diseases and vary with farming practices in the Mediterranean Thau lagoon, France. Aquac. Environ. Interact. 2, 215–237. doi:10.3354/aei00041.

Pernet, F., Lupo, C., Bacher, C., and Whittington, R. J. (2016). Infectious diseases in oyster aquaculture require a new integrated approach. Philos. Trans. R. Soc. B Biol. Sci. 371, 20150213. doi:10.1098/rstb.2015.0213.

Pernet, F., Tamayo, D., Fuhrmann, M., and Petton, B. (2019). Deciphering the effect of food availability, growth and host condition on disease susceptibility in a marine invertebrate. J. Exp. Biol. 222, jeb210534. doi:10.1242/jeb.210534.

Petton, B., Bruto, M., James, A., Labreuche, Y., Alunno-Bruscia, M., and Le Roux, F. (2015). *Crassostrea gigas* mortality in France: the usual suspect, a herpes virus, may not be the killer in this polymicrobial opportunistic disease. Front. Microbiol. 6, 686. doi:10.3389/fmicb.2015.00686.

Petton, B., Pernet, F., Robert, R., and Boudry, P. (2013). Temperature influence on pathogen transmission and subsequent mortalities in juvenile Pacific oysters *Crassostrea gigas*. Aquac. Environ. Interact. 3, 257–273. doi:10.3354/aei00070.

Pfaffl, M. W. (2001). A new mathematical model for relative quantification in real-time RT-PCR. Nucleic Acids Res. 29, 45e–445. doi:10.1093/nar/29.9.e45.

R: a language and environment for statistical computing, 2008; R Development Core Team, R Foundation for Statistical Computing, Vienna, Austria [http://www.R-project.org].

Renault, T., and Cochennec, N. (1995). Chlamydia-like organisms in ctenidia and mantle cells of the Japanese oyster *Crassostrea gigas* from the French Atlantic coast. Dis. Aquat. Organ. 23, 153–159.

Rico-Villa, B., Pouvreau, S., and Robert, R. (2009). Influence of food density and temperature on ingestion, growth and settlement of Pacific oyster larvae, *Crassostrea gigas*. Aquaculture 287, 395–401. doi:10.1016/j.aquaculture.2008.10.054.

Rognes, T., Flouri, T., Nichols, B., Quince, C., and Mahé, F. (2016). VSEARCH: a versatile open source tool for metagenomics. PeerJ 4, e2584. doi:10.7717/peerj.2584.

Rubio, T., Oyanedel, D., Labreuche, Y., Toulza, E., Luo, X., Bruto, M., et al. (2019). Species-specific mechanisms of cytotoxicity toward immune cells determine the successful outcome of *Vibrio* infections. Proc. Natl. Acad. Sci. 116, 14238–14247. doi:10.1073/pnas.1905747116.

Rungrassamee, W., Klanchui, A., Maibunkaew, S., and Karoonuthaisiri, N. (2016). Bacterial dynamics in intestines of the black tiger shrimp and the Pacific white shrimp during *Vibrio harveyi* exposure. J. Invertebr. Pathol. 133, 12–19. doi:10.1016/j.jip.2015.11.004.

Samain, J. F., Dégremont, L., Soletchnik, P., Haure, J., Bédier, E., Ropert, M., et al. (2007). Genetically based resistance to summer mortality in the Pacific oyster (*Crassostrea gigas*) and its relationship with physiological, immunological characteristics and infection processes. Aquaculture 268, 227–243. doi:10.1016/j.aquaculture.2007.04.044.

Takenaka, S., Yoshikawa, T., Kadowaki, S., Okunishi, S., and Maeda, H. (2017). Microflora in the soft tissue of the Pacific oyster *Crassostrea gigas* exposed to the harmful microalga *Heterosigma akashiwo*. Biocontrol Sci. 22, 79–87. doi:10.4265/bio.22.79.

Travers, M.-A., Mersni Achour, R., Haffner, P., Tourbiez, D., Cassone, A.-L., Morga, B., et al. (2014). First description of French *V. tubiashii* strains pathogenic to mollusk: I. Characterization of isolates and detection during mortality events. J. Invertebr. Pathol. 123, 38–48. doi:10.1016/j.jip.2014.04.009.

Turner, J. W., Good, B., Cole, D., and Lipp, E. K. (2009). Plankton composition and environmental factors contribute to *Vibrio* seasonality. ISME J. 3, 1082–1092.

Wang, A., Ran, C., Wang, Y., Zhang, Z., Ding, Q., Yang, Y., et al. (2019). Use of probiotics in aquaculture of China—a review of the past decade. Fish Shellfish Immunol. 86, 734–755. doi:10.1016/j.fsi.2018.12.026.

Wendling, C. C., Fabritzek, A. G., and Wegner, K. M. (2017). Population-specific genotype x genotype x environment interactions in bacterial disease of early life stages of Pacific oyster larvae. Evol. Appl. 10, 338–347. doi:10.1111/eva.12452.

Wilsey, B. J., and Polley, H. W. (2002). Reductions in grassland species evenness increase dicot seedling invasion and spittle bug infestation. Ecol. Lett. 5, 676–684.

Wilsey, B. J., and Potvin, C. (2000). Biodiversity and ecosystem functioning: importance of species evenness in an old field. Ecology 81, 887–892.

Wittebolle, L., Marzorati, M., Clement, L., Balloi, A., Daffonchio, D., Heylen, K., et al. (2009). Initial community evenness favours functionality under selective stress. Nature 458, 623–626. doi:10.1038/nature07840.

Woodhams, D. C., Vredenburg, V. T., Simon, M.-A., Billheimer, D., Shakhtour, B., Shyr, Y., et al. (2007). Symbiotic bacteria contribute to innate immune defenses of the threatened mountain yellow-legged frog, *Rana muscosa*. Biol. Conserv. 138, 390–398. doi:10.1016/j.biocon.2007.05.004.

Zaneveld, J. R., McMinds, R., and Vega Thurber, R. (2017). Stress and stability: applying the Anna Karenina principle to animal microbiomes. Nat. Microbiol. 2, 17121. doi:10.1038/nmicrobiol.2017.121.

Zhang, Y., Chen, H. Y. H., and Reich, P. B. (2012). Forest productivity increases with evenness, species richness and trait variation: a global meta-analysis. J. Ecol. 100, 742–749. doi:10.1111/j.1365-2745.2011.01944.x.

