## Supplementary material for "Microbiota composition and evenness predict survival rate of oysters confronted to Pacific Oyster Mortality Syndrome": Legends of Supplementary Files

**Legends of Supplementary Material**

Clerissi et al.

**Legends of supplementary figures**

**Supplementary Figure 1. Rarefaction analyses of species richness for oyster microbiota.**

**Supplementary Figure 2. Heatmap of bacterial genera that were significantly different between resistant and susceptible oyster families in the controlled condition (hatchery).** Only bacterial genera with a frequency above 4% in at least one sample are shown. Frequencies above and below 4% are displayed in red and blue, respectively.

**Supplementary Figure 3. Heatmap of bacterial genera that were significantly different between resistant and susceptible oyster families in the infectious condition (field).** Only bacterial genera with a frequency above 4% in at least one sample are shown. Frequencies above and below 4% are displayed in red and blue, respectively.

**Legends of supplementary tables**

**Supplementary Table 1. Number of sequences and OTUs.**

**Supplementary Table 2. OTU annotations and abundances in the 83 oyster microbiota.**

**Supplementary Table 3. Metadata and alpha diversity indices.**

**Supplementary Table 4. Significant bacterial genera.** For *DESeq* columns, *resistant* and *susceptible* indicate if abundances of each bacterial genus were higher in resistant or susceptible oyster families, respectively. NS: not significant. NT: not tested (absent from the dataset).

**Supplementary Table 5. Significant bacterial families.** For *DESeq* columns, *resistant* and *susceptible* indicate if abundances of each bacterial family were higher in resistant or susceptible oyster families, respectively. NS: not significant. NT: not tested (absent from the dataset).
