## Supplementary figures and images for "Microbiota composition and evenness predict survival rate of oysters confronted to Pacific Oyster Mortality Syndrome"

### Supplementary Figure 1

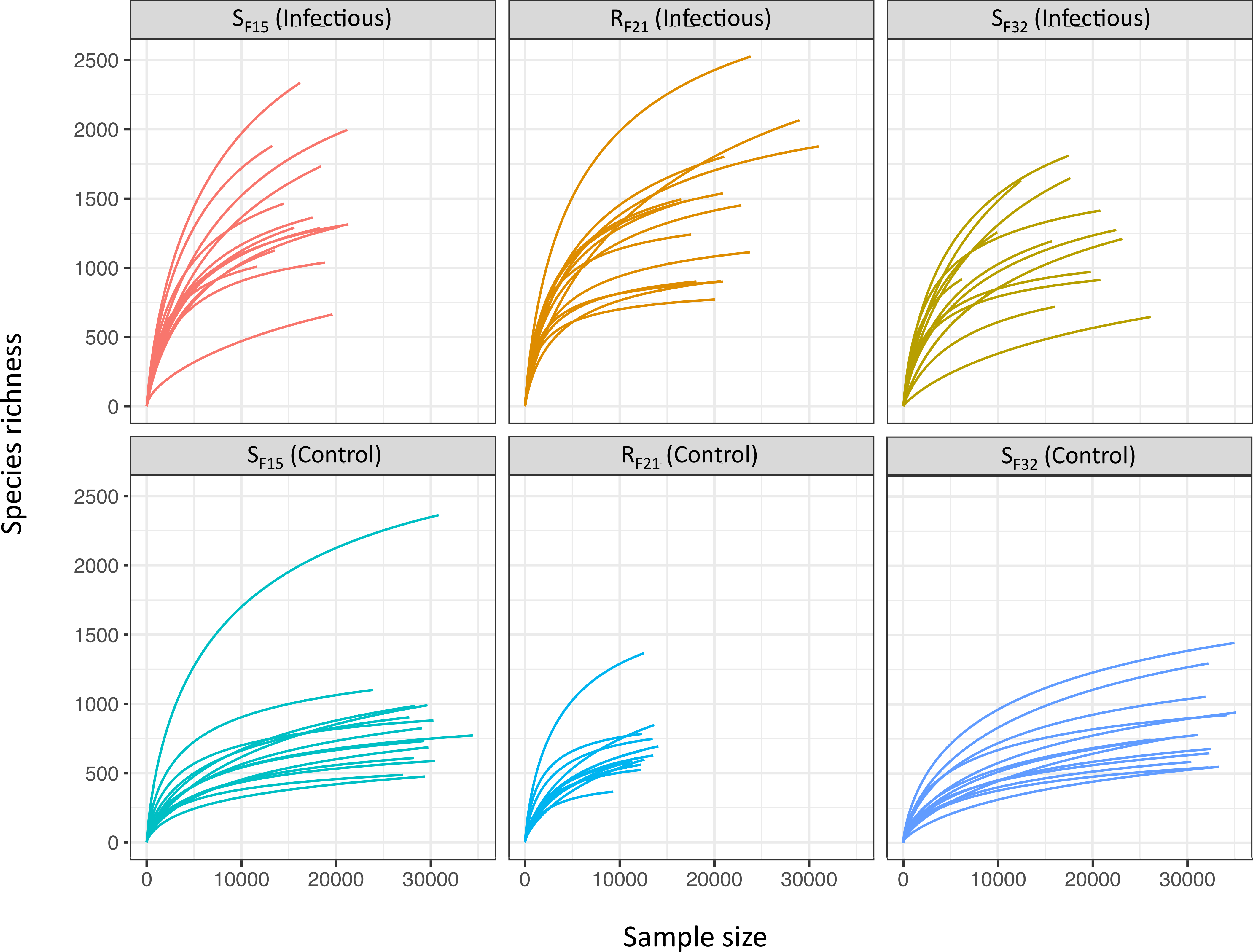

### Supplementary Figure 2

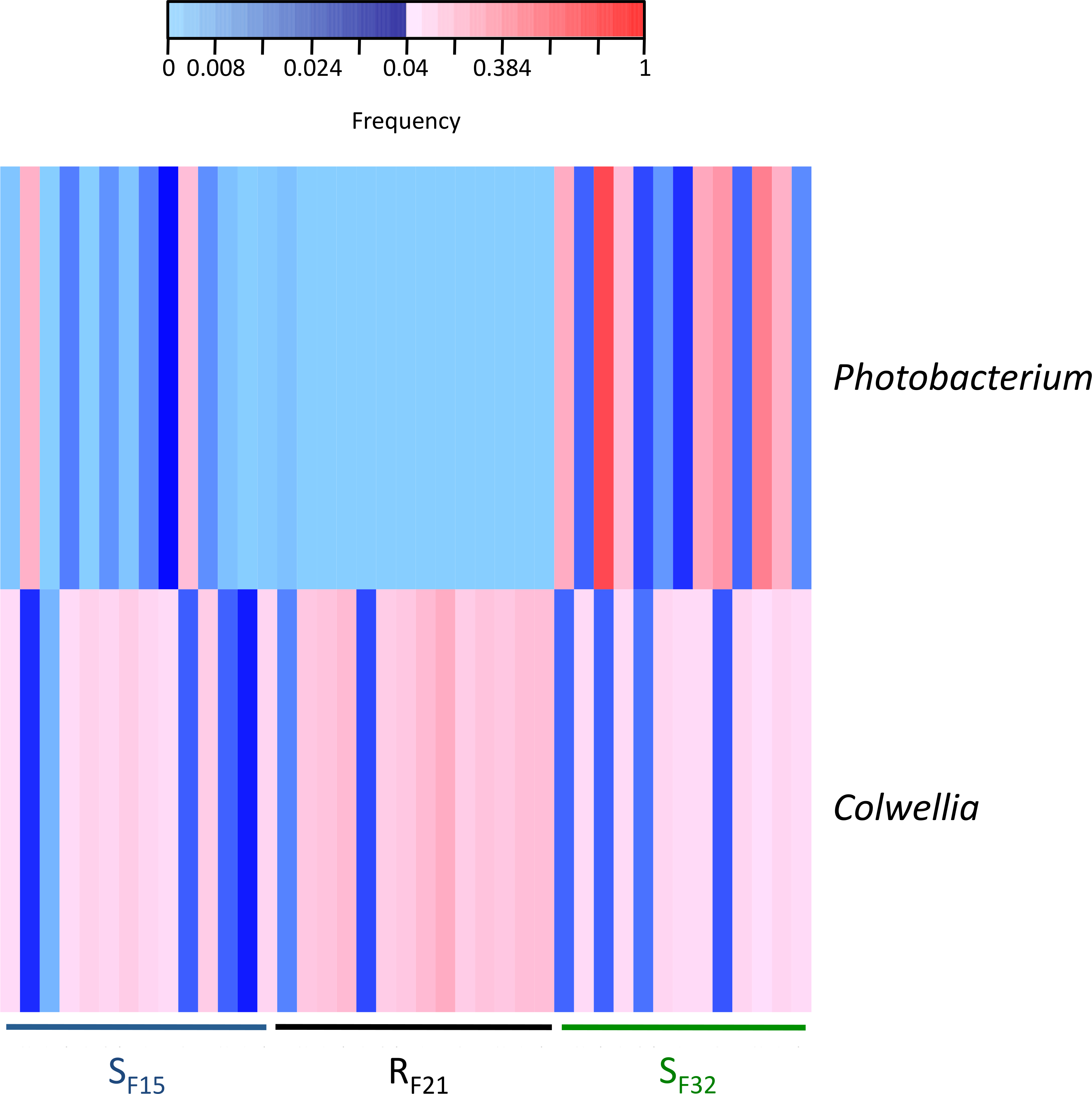

### Supplementary Figure 3

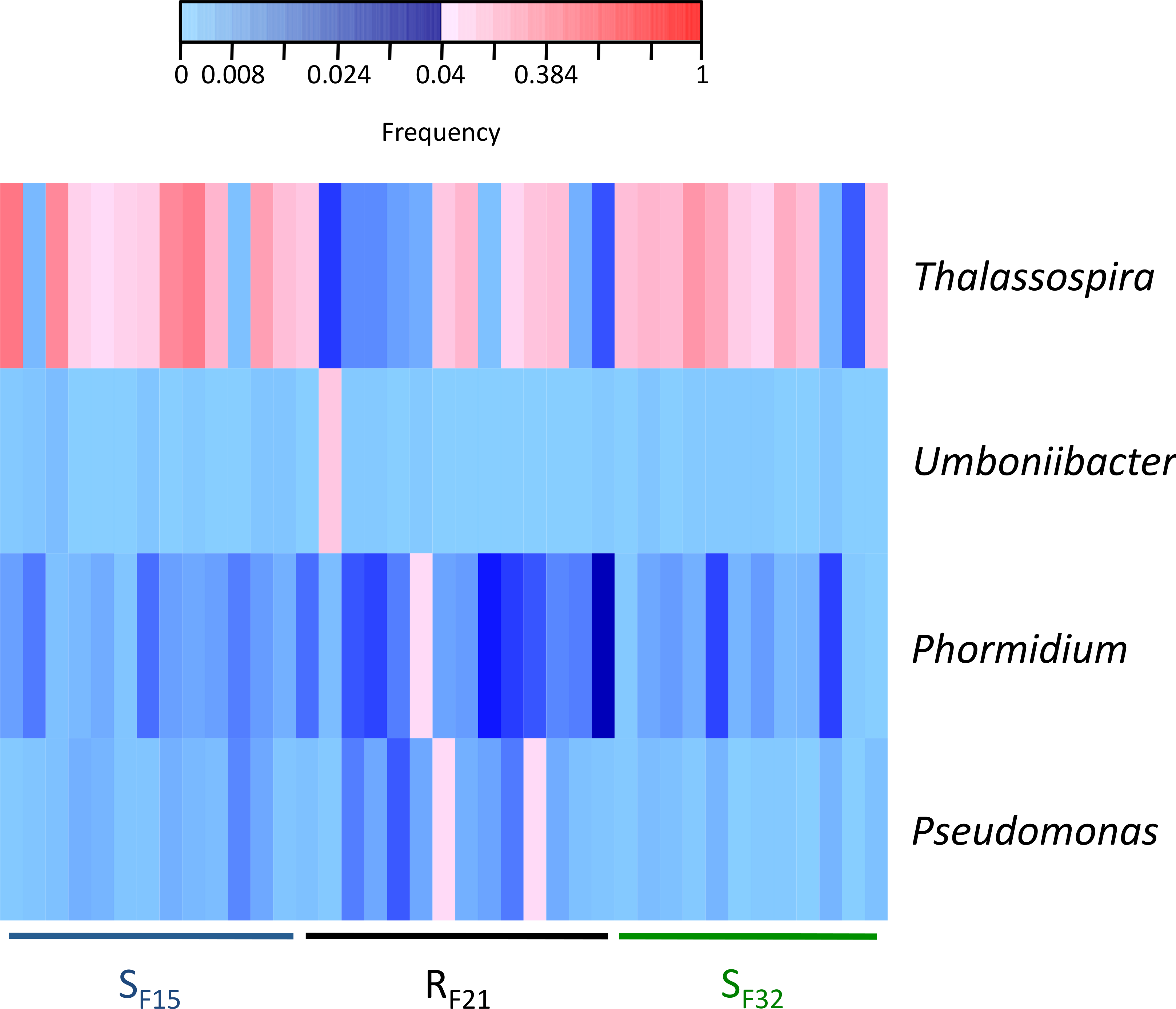
